# Genoprotective effects of Chaga mushroom (*Inonotus obliquus*) polysaccharides in UVB-exposed embryonic zebrafish (*Danio rerio*) through coordinated expression of DNA repair genes

**DOI:** 10.1101/2020.06.19.161182

**Authors:** Jehane Ibrahim Eid, Swabhiman Mohanty, Biswadeep Das

## Abstract

**Background:** Chaga mushroom *(Inonotus obliquus)* is one of the most promising antioxidants with incredible health-promoting effects. Chaga polysaccharides (IOP) have been reported to enhance immune response and alleviate oxidative stress during development. However, the effects of IOP on the genotoxicity in model organisms are yet to be clarified.

**Methods:** Zebrafish embryos (12 hours post fertilization, hpf) were exposed to transient UVB (12 J/m^2^/s, 310 nm) for 10 secs using a UV hybridisation chamber, followed by IOP treatment (2.5 mg/mL) at 24 hpf for up to 7 days post fertilization (dpf). The genotoxic effects were assessed using acridine orange staining, alkaline comet assay, and qRT-PCR for screening DNA repair genes.

**Results:** We found significant reduction in DNA damage and amelioration of the deformed structures in the IOP-treated zebrafish exposed to UVB (p < 0.05) at 5 dpf and thereafter. In addition, the relative mRNA expressions of *XRCC-5, XRCC-6, RAD51, P53,* and *GADD45* were significantly upregulated in the IOP-treated UVB-exposed zebrafish. Pathway analysis demonstrated coordinated regulation of DNA repair genes, suggesting collective response during UVB exposure.

**Conclusions:** Overall, IOP treatment ameliorated the genotoxic effects in UVB-exposed zebrafish embryos, which eventually assisted in normal development. The study suggested the efficacy of Chaga mushroom polysaccharides in mitigating UV-induced DNA damage.

## Introduction

The National Cancer Institute (NCI), United States has recently intensified its focus on natural products such as plants, marine organisms, and certain microorganisms for use in drug and biosimilar discovery [1] In this regard, Chaga *(Inonotus obliquus)* mushroom is one such biologically relevant product, which has gained increasing attention worldwide because of its high nutritional and medicinal values, such as antioxidant, anti-inflammatory, immune booster, antidiabetic, antiviral and several others [2–5]. In particular, *Inonotus obliquus* polysaccharide (IOP) represents the most active ingredient of Chaga mushroom, known for treating and preventing various diseases such as tumors, metabolic disorders, and many chronic diseases in folk medicine. Such powerful beneficial effects are largely due to the immunomodulatory, antiinflammatory and anti-oxidative properties of *Inonotus obliquus* polysaccharide. Chaga tea is popular and consumed in many countries for several remedies and health promoting effects [5,6].

Because DNA mutations are considered as the prime reason for developing cancer and genetic disorders, it is important to maintain the integrity of DNA in the cell for proper functioning. Large-scale DNA damage can be deleterious to the cell and are caused mainly by alkylating agents (transform a functional base into a mutagenic one), hydrolytic deamination (lead to base alterations), dyes (ethidium bromide), reactive oxygen species (intrinsic) and ionizing radiations (ultraviolet light, UV) that render maximum DNA damage [7–9]. UV light comprises three broad categories based on their wavelength: UVA (320–400 nm), UVB (290–320 nm), and UVC (240–290 nm). UVC is the most harmful to living organism due to its exorbitantly high frequency and is mostly absorbed by the stratospheric ozone [10,11]. A part of UVB radiation is also absorbed in the stratosphere, though majority of UVB radiation reaches the earth’s surface, and is responsible for serious health comes, such as skin cancer by inducing DNA damage. UVB results in two major types of mutagenic DNA lesions: cyclobutane–pyrimidine dimers, and pyrimidine adducts 6–4 photoproducts, in addition to their Dewar valence isomers [11].

Although there are in vivo repair mechanisms to manage and correct DNA mutations and maintain is integrity such as photo-reactivation, base/nucleotide excision repair, DNA replication related methyl mismatch repair, in addition to large scale and nonspecific repair systems (SOS response and apoptosis), the degree of repair depends on the extent of damage and the status of the repair systems of the human body [10]. Therefore, intake of natural antioxidants and immune boosters are highly recommended for maintaining a healthy state of the DNA and eventually the whole body [4,6]. In this regard, Chaga mushroom is a promising supplement whose benefits have been assessed in several metabolic and chronic disorders. However, it is imperative to understand the effect of Chaga mushroom on the genotoxic profile in vivo to understand if it has any effect on DNA damage reversal.

Recently, many international toxicity researches have utilized zebrafish (Danio rerio) as an in vivo model organism because it possesses several measurable indicators in ecotoxicology, such as small size, high fecundity, well-characterized embryonic ontogenesis, transparent embryos, and rapid development [12]. Besides, zebrafish has more than 70% genome similarity with the humans that render it to effectively model any human disease with high phenotypic similarity and convenience [13]. Zebrafish is an ideal model for assessing DNA damage and its counter mechanisms because zebrafish genomic DNA contains DNA repair genes orthologues that participate in DNA repair mechanisms [12,13], as well as zebrafish DNA is amenable to genetic manipulation using morpholinos, shRNA, or Crisprs to assesses the role of genes during DNA repair response [14]. Zebrafish has been used as in vivo model organism for testing a plethora of exogenous agents comprising drugs, nanoparticles, organic and inorganic compounds, besides being exposed to different stress environments. UV-induced zebrafish models have also been developed to study the expression of DNA repair genes and tumor suppressor genes regulation [15–17]. Furthermore, transparent zebrafish casper mutants have been consistently used for tumour-related and toxicological studies for assessing the phenotypic manifestations during development [18]. Such avenues will be worthy to explore the specific DNA repair genes that might play additional roles in embryological development. The objective of this study was to assess the molecular mechanism of IOP in UVB-exposed zebrafish during early development.

## Results

### Development analysis upon IOP exposure in zebrafish embryos

Embryo development varied across the three groups; the UVB-exposed zebrafish showed severe structural aberrations, such as yolk sac edema, loss of vital structures, and slow development compared to control and IOP treated UVB-exposed groups (Fig 1). Average mortality was high in the UVB-exposed group (80%) during the course of development compared to IOP-treated (15%) and control (10%). Interestingly, IOP-treated UVB exposed group showed structural deformations like pericardial edema and spinal curvature in the early days of development (3 dpf), which were significantly ameliorated during late development (≥ 5 dpf). The embryos remained healthy without showing any signs of mortality at the later stages of development in the IOP-treated group. Most vital statistics such as development time, hatching rate and heart rate was similar in the control and IOP-treated UVB exposed group, which indicated that IOP promoted the development of zebrafish embryos as they would have developed under normal conditions.

**Fig. 1.**
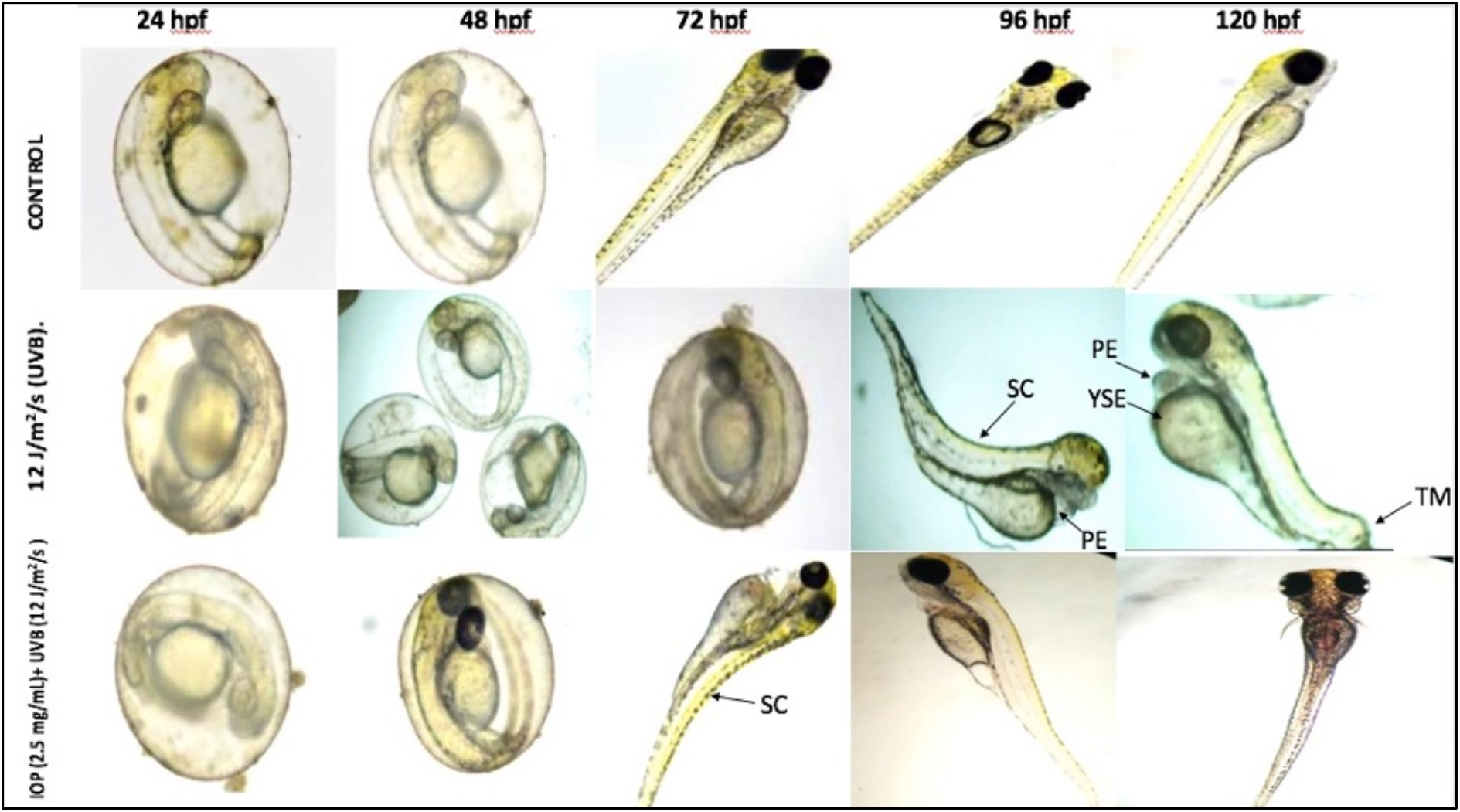
Bright field images of morphological analysis of zebrafish embryos (1 dpf to 4 dpf) in the three different zebrafish groups (control, UVB-exposed, IOP treated UVB-exposed).

### Acridine orange staining

The level of DNA damage was assessed by the interaction of DNA binding AO dye, which was observed to be significantly uptaken by the UVB-exposed zebrafish embryos (5 dpf) compared to control and IOP-treated UVB-exposed groups. The uptake was visualized as intense green fluorescence (Fig 2A), which was graphically represented as a histogram (Fig 2B). Moreover, IOP-treated UVB-exposed group showed remarkably low green fluorescence demonstrating less DNA interactions with the dye. The results showed that IOP assisted in maintaining the integrity of the DNA, owing to less penetration of AO into the nucleus, which was similar to that in control embryos. These results indicated significantly increased apoptosis (p < 0.05) in the UV-exposed group without IOP exposure compared to IOP-treated UVB exposed group, which implied that IOP conferred protection against DNA damage.

**Fig. 2.**
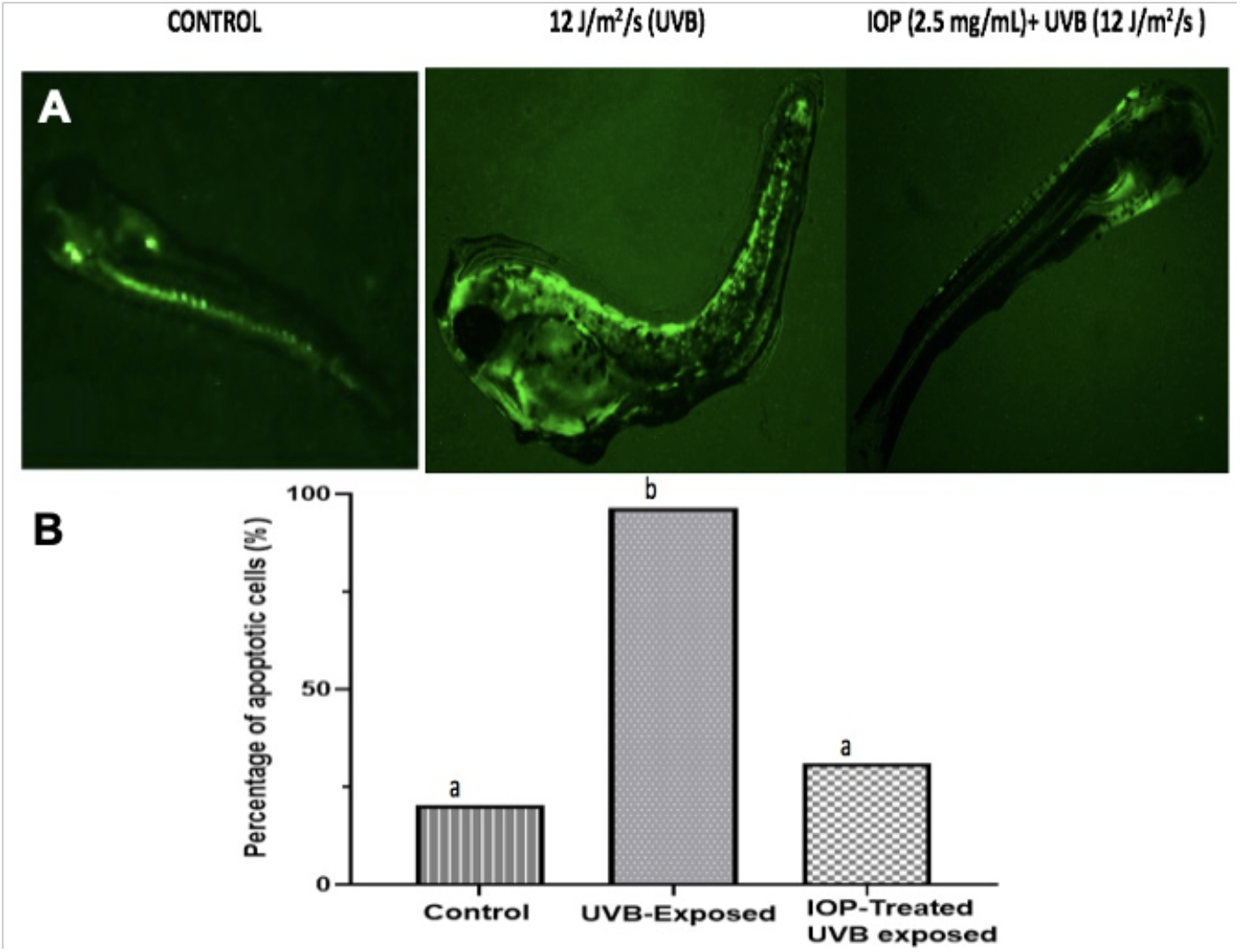
A) Acridine orange staining of zebrafish embryos (5 dpf) for the three zebrafish (groups control, UVB-exposed, IOP treated UVB-exposed), showing intensive green fluorescence as significant uptake of AO dye due to cellular-compartment and DNA damage. B) shows the histogram generated by Image J analysis showing the percentage of apoptotic cells in the three groups.

### Genotoxicity assessment using alkaline comet assay

Genotoxic assessment using the alkaline comet assay by analyzing the tail intensity in the three zebrafish groups revealed varied genotoxic effects, exhibiting distinct comet heads and tail regions (Fig 3). On 5 dpf, the tail intensity was significantly increased in the UVB-exposed zebrafish embryos by almost three-fold and four-fold compared to IOP-induced UVB exposed and control groups, respectively (Fig 4). On 7dpf, the tail intensity was further significantly increased in the UVB-exposed zebrafish embryos (86.6 %) by almost four-fold compared to both IOP-induced UVB exposed (32.1%) and control groups (26.1%), respectively (Fig 4). These results indicated that IOP acted as an anti-genotoxic compound and assisted in the amelioration of genotoxicity caused due to UVB damage, thereby allowing the normal development of the zebrafish embryos.

**Fig. 3.**
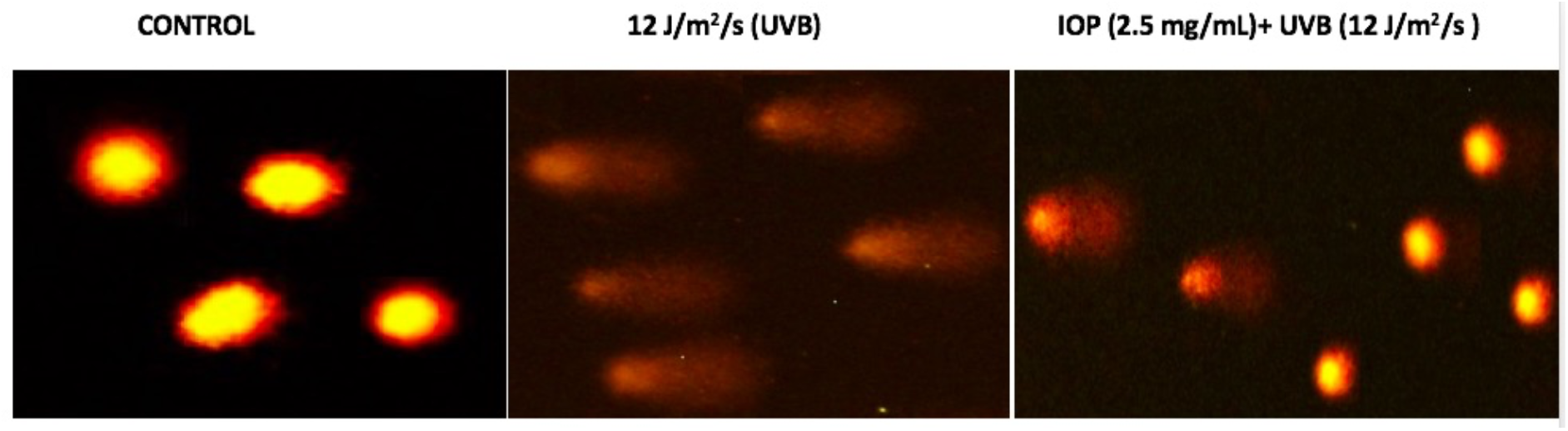
Alkaline comet assay showing distinct comet head and tail after ethidium bromide staining and fluorescent microscopy in the three zebrafish (groups control, UVB-exposed, IOP treated UVB-exposed) at 5dpf.

**Fig. 4.**
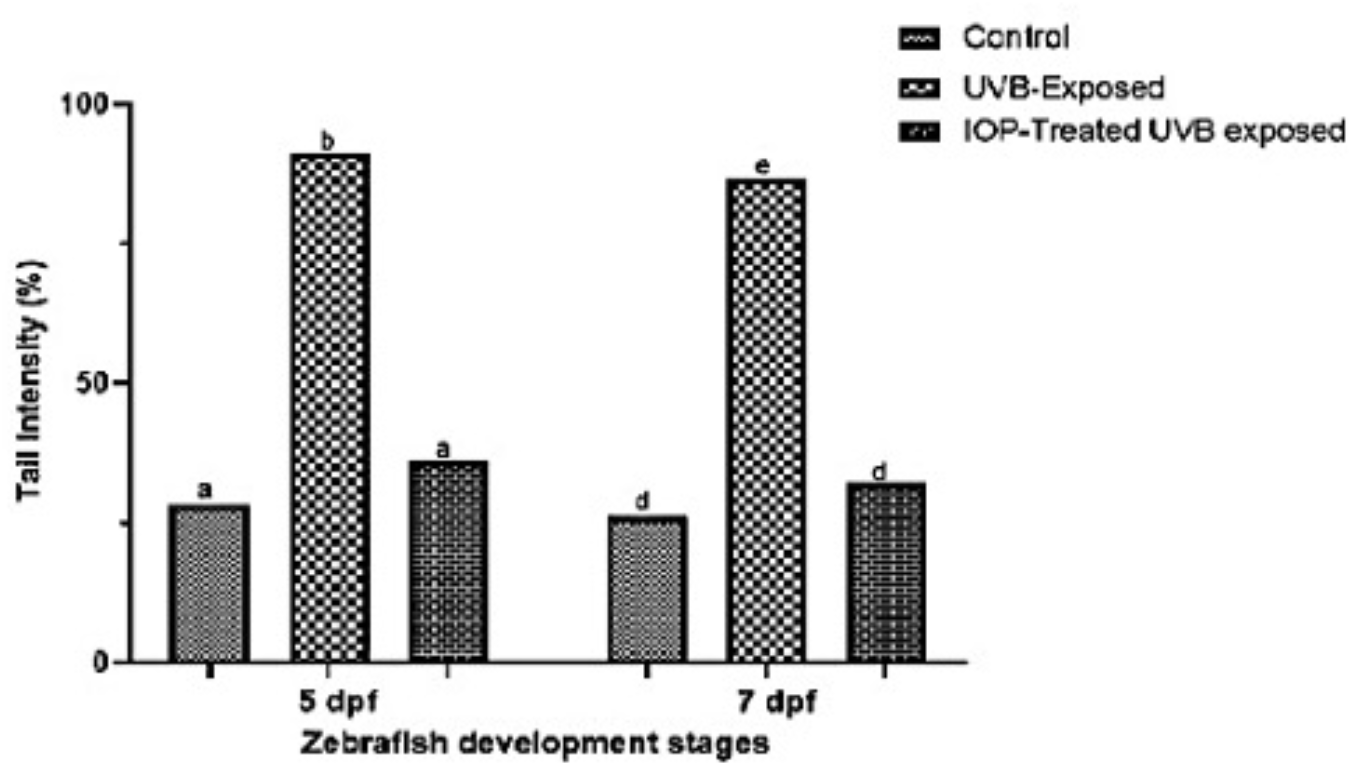
Comet assay histogram at 5 dpf and 7 pdf generated by analyzing the tail intensity (DNA fragmentation) using Image J in the three zebrafish (groups control, UVB-exposed, IOP treated UVB-exposed) at 5dpf.

### Gene expression analysis upon IOP exposure in zebrafish embryos

qRT-PCR analysis showed that the relative expression of DNA repair genes *XRCC5, XRCC6, RAD51, GADD45, P53,* and *BAX* was significantly up-regulated in the IOP-treated UVB-induced zebrafish compared to UVB-exposed and control groups (p < 0.05). The mean expression of *XRCC5, XRCC6, RAD51, GADD45,* and *P53* in the IOP-treated UV-exposed zebrafish was significantly increased by 1.87, 1.73, 2.41, 2.55, and 1.65 times in comparison to control fish. In the UVB-exposed group, the mean expression of all the above-mentioned genes, except *BAX* was lower than or equal to control zebrafish (Fig. 5), indicating that UVB exposure induced substantial damage to the DNA repair system. The increased expression of *BAX* gene in the UVB-exposed group could attribute to the increased apoptosis, leading to rapid cell death. These findings demonstrated that IOP enhanced the expression of DNA repair genes upon UVB exposure, thereby promoting DNA protective effects and eventually assisting in the development of zebrafish.

**Fig. 5.**
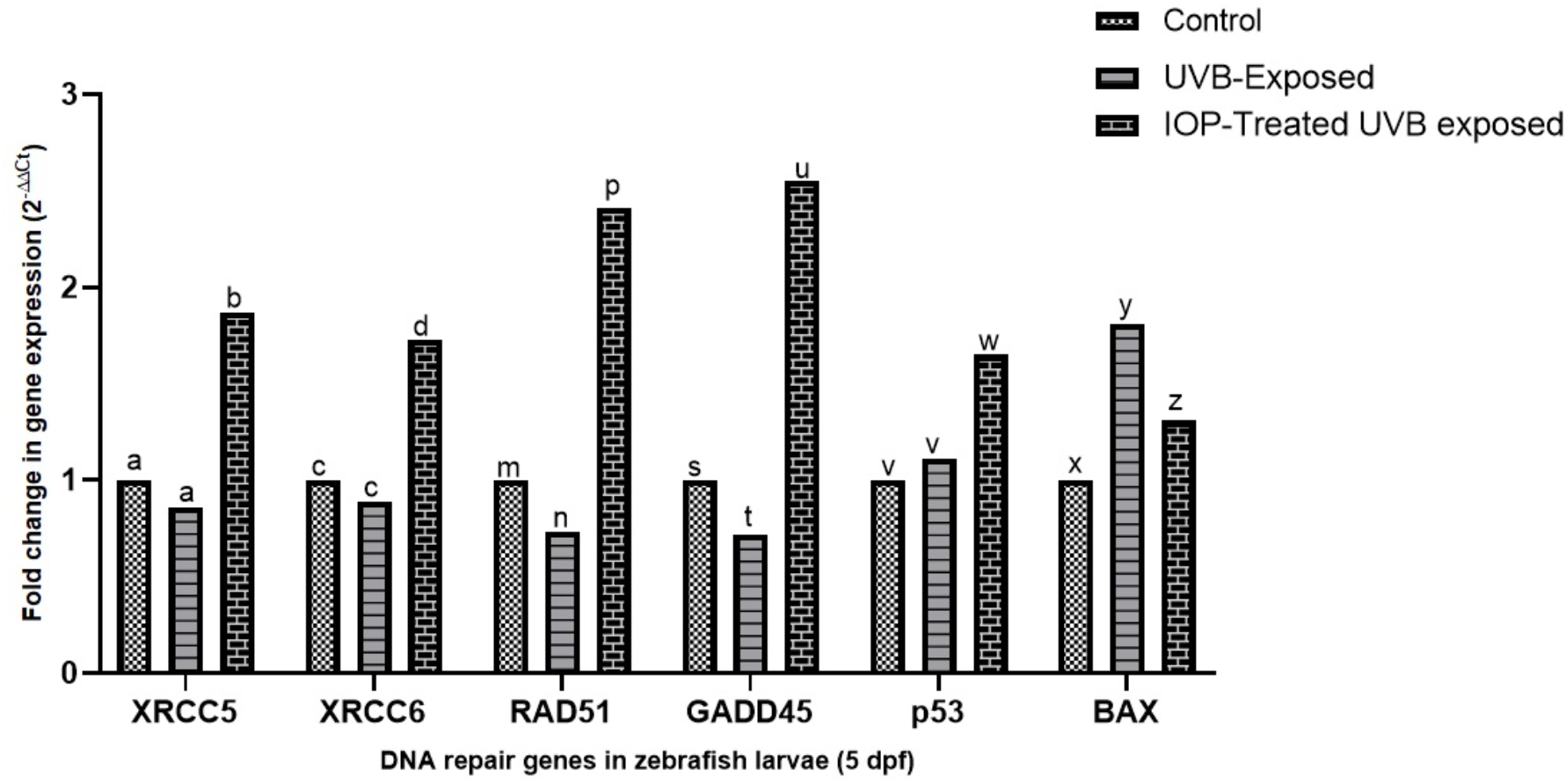
qRT-PCR bar graphs showing the fold change in the gene expression in UVB-exposed and IOP-treated UVB exposed groups compared to that of control.

## Discussion

Mushrooms possessing medicinal values, such as Ganoderma, Chaga, and many others are currently being explored significantly because of their tremendous benefits without inducing any undesirable effects, besides possessing beneficial attributes such as anti-inflammatory, anti-oxidant, and anti-tumor [12–17]. Given the wide benefits of Chaga mushroom, the effects of Chaga mushroom on the genotoxicity profile of an organism will be intriguing to explore. In this study, we found that hot water extracted-Chaga polysaccharides significantly mitigated genotoxicity by enhancing the expression of DNA repair genes in UVB-induced zebrafish embryos. The major polysaccharide found in Chaga mushroom was β-glucan, a polymer of β-D glucose, which is documented to possess several health benefits [18]. β-glucan has been reported to have anti-diabetic, anti-proliferative properties, and anti-tumorigenic properties [19]. Such beneficial effects could be attributed to the ability of β-glucan to scavenge free radicals generated due to endogenous or exogenous agents, such as ionizing UVB radiation, thereby assisting the cells to repair DNA [20]. Several studies have reported that β-glucan readily facilitates the reduction of both simple and complex chromosomal aberrations, and confer protection to the DNA against single-strand and double-strand breaks [21], in addition to enhancing the DNA repair system [22]. β-glucan thus confers protection to DNA through its antioxidant activity and by enhancing the DNA repair system, besides being water soluble that allows it to be readily uptaken by the cell without any perturbations in the cellular processes. Recently, several bioactive polysaccharides isolated from natural sources have been given much attention in clinical pharmacology [23]. Such polysaccharides can be modified by chemical methods to improve the antitumor activity of polysaccharides and their clinical qualities, such as water solubility. Therefore, IOP could be regarded as a potential adjunct along with conventional chemotherapy for several ailments and can have wide application in health clinics [23,24].

The present study revealed that continuous exposure to IOP reversed the structural deformations that were induced due to UVB exposure in zebrafish embryos. Severe organ damage, and loss of vital structures that occurred in the UVB-exposed zebrafish were significantly ameliorated in the IOP-treated UVB induced groups. This suggested that IOP conferred protection to the zebrafish embryos against exogenous agents. During early development, DNA damage results in severe structural deformations, which was evidently seen in the UVB-exposed zebrafish. The reversal of such damage and eventual normal development of the zebrafish embryos could be solely attributed to continuous exposure to IOP that can be further explained due to the presence of high amount of beta-glucans [24–27]. Moreover, we observed the reversal of DNA damage by comet assay; and DNA fragmentation, which was measured as the comet tail intensity was significantly reduced in the IOP-treated UVB-exposed group. This suggested that IOP was anti-genotoxic and conferred protection to DNA by maintaining its integrity. Furthermore, IOP also reduced the number of apoptotic cells in the zebrafish embryos as demonstrated by acridine orange staining, whereas the apoptosis was significantly increased in the UVB-exposed zebrafish. This could also be correlated with increased expression of the Bcl2 associated X gene, *Bax* that promoted apoptosis in the UVB-exposed zebrafish. However, the *Bax* gene was downregulated in IOP-treated group compared to UVB exposed group, which suggested that IOP was inhibited apoptosis and increased cellular longevity.

Continuous exposure to IOP significantly reduced the amount of damaged DNA and thus, the embryos developed normally [28]. Such DNA protective effects could be explained by the enhanced expression of several DNA repair genes. In this study, we observed significantly elevated expression of *xrcc5, xrcc6, rad51,* and *gadd45* genes in the IOP-treated UVB-exposed group compared to UVB-exposed group. *Xrcc5* and *xrcc6* encode Ku70 and Ku80 DNA repair proteins that function collectively during non-homologous end joining during DNA damage.

Overexpression of these genes have also been reported in several disorders involving DNA damage to correct the DNA [29,30]. *Rad51* encodes a protective enzyme that facilitates the repair of double strand DNA breaks by catalyzing strand transfer between a broken sequence and its undamaged homologue to allow re-synthesis of the damaged region [31,32]. It has a major role in homologous recombination, and assists in recombination repair during extensive DNA damage [32]. The substantial increase in the expression of these genes indicated prompt recruitment of DNA repair proteins and processing of damaged DNA for the correction of damaged DNA, in addition to maintaining its integrity. We also found increased expression of *GADD45* gene, which encodes the growth arrest and DNA damage proteins that have a critical role during embryogenesis by regulating differentiation (by inducing the zygotic gene expression) along with protecting DNA from spontaneous damage, mainly by efficient recognition and repair of spontaneous DNA damages by ten-eleven translocation methylcytosine dioxygenase 1 (TET)-mediated DNA demethylation [33–36]. This gene was the most highly expressed in the IOP-treated UVB-exposed group, which suggested that IOP greatly enhanced the expression of *GADD45* proteins, which eventually assisted the embryos in normal development. Network analysis showed that all the DNA repair genes encoding proteins acted in a coordinate manner during cellular signaling against DNA damage. Furthermore, *tp53* acted as the central player regulating the coordinate response of all the genes involved in DNA repair (Fig 7 6). These findings demonstrated that all the genes need to act in a coordinate manner during DNA repair under the control of *tp53.*

**Fig. 6.**
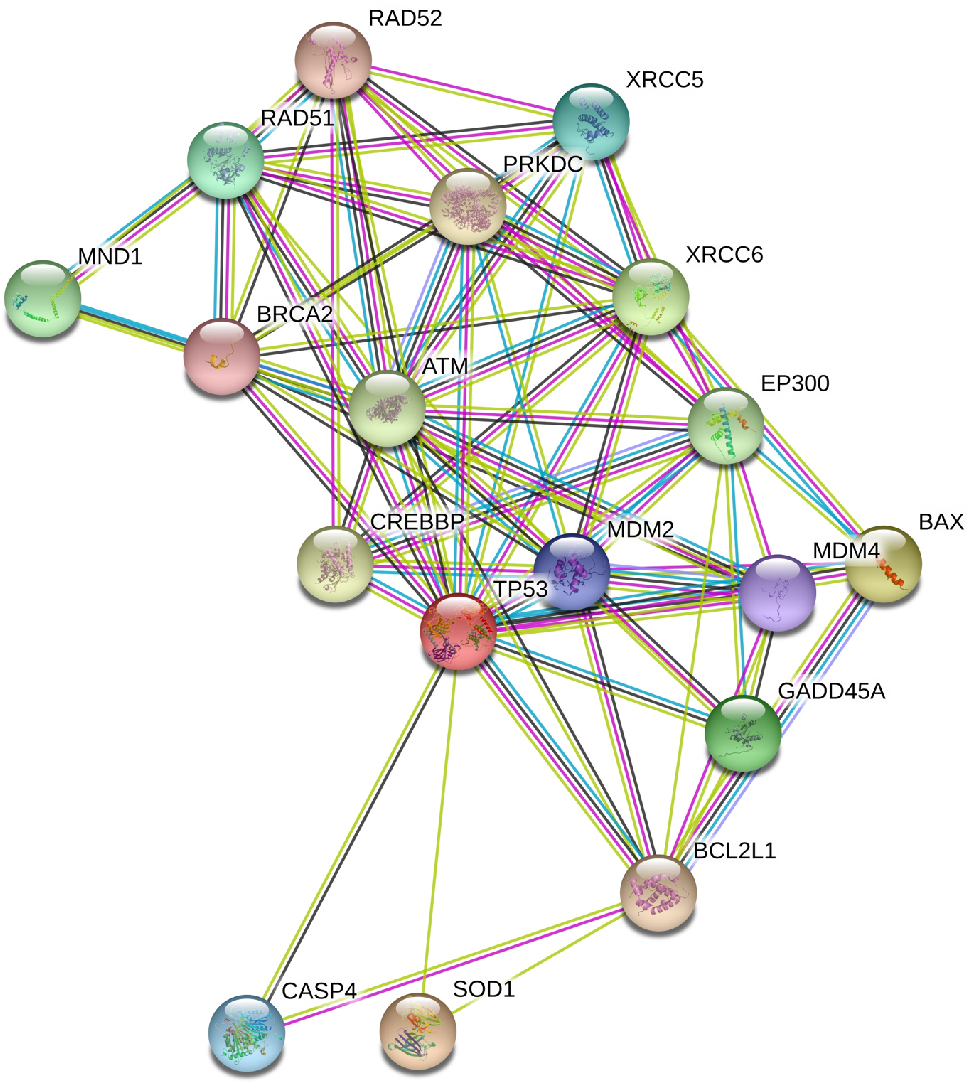
Network analysis using String software demonstrating the coordinated association of the DNA repair genes involved during UV induced DNA repair.

**Fig. 7.**
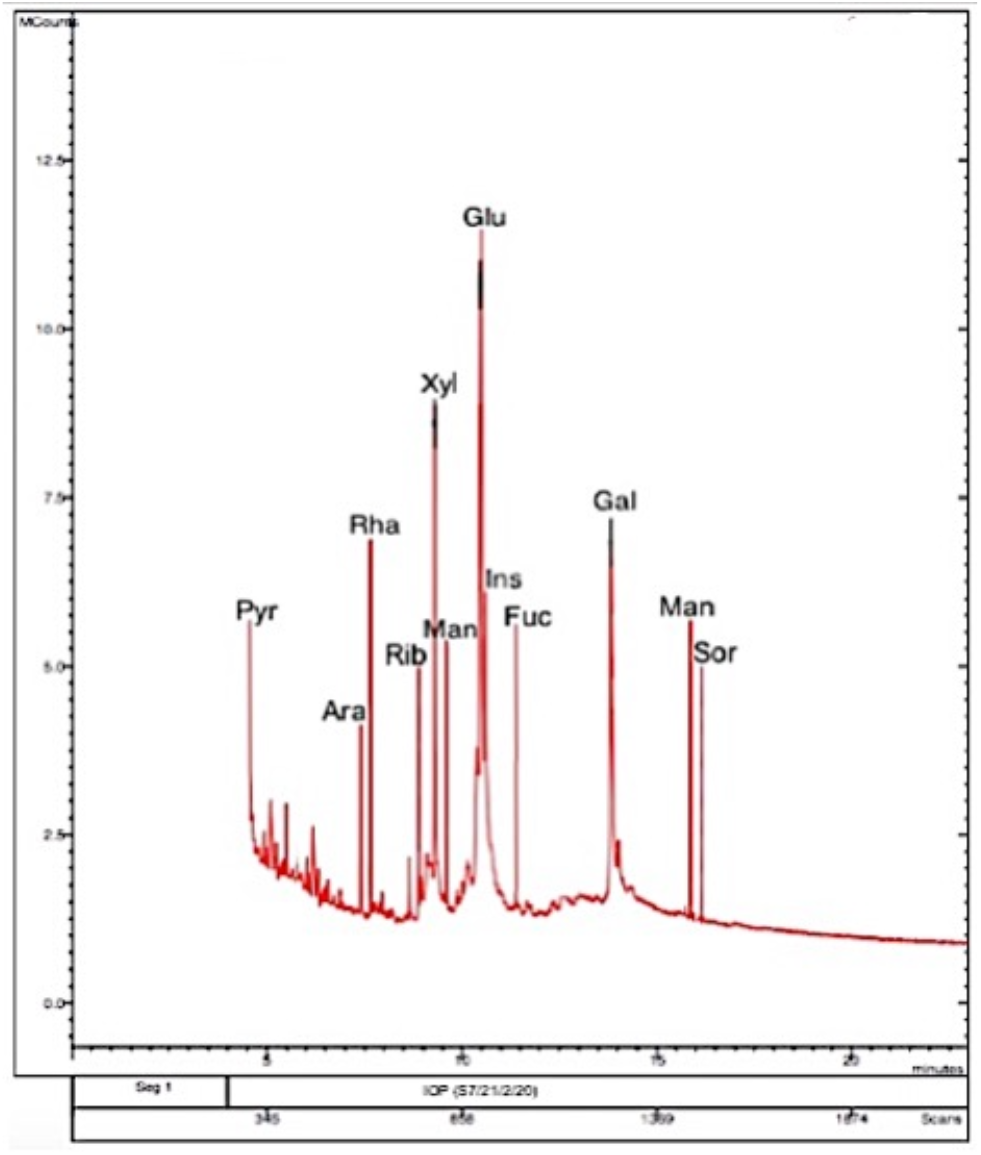
GC-MS chromatogram depicting the retention time peaks for different monomers of hot water-extracted Chaga mushroom polysaccharides.

In vivo developmental analyses showed that continuous exposure to IOP extract did not exhibit any significant developmental deformities in embryonic zebrafish, which developed in line with control embryos. This finding suggested that Chaga mushroom aided in normal development of the zebrafish and reduced DNA damage in the developing embryos.

Therefore, Chaga mushroom could be a potent natural therapeutic for ameliorating disorders linked to DNA damage.

## Methods

### Extraction of Chaga mushroom polysaccharides and GC-MSMS analysis

Extraction of Chaga mushroom polysaccharides (IOP) was performed using hot water-ethanolic extraction method according to Eid et al., 2020 [12]. Briefly, 10 g of Siberian grade Chaga chunks were dried, ground to fine powder and dissolved in 150 mL followed by refluxing at 70 °C, and vacuum dried and concentrated using 3 volumes of 95% ethanol. The solution was then centrifuged at 5000 rpm for 10 min, and the supernatant was dried and treated with Sevag reagent (chloroform:butanol in the ratio 4:1) to remove the proteins. The solution was then oven-dried and mixed with distilled water (5 g in 250 mL w/v), followed by chromatographic analysis and spectrophotometric confirmation [12].

GC-MS analysis was performed using trimethylsilylation reagent, TMS (N-Trimethylsilylimidazole) as per previous protocol [12]. The area normalization method was employed to estimate the molar ratio of the monosaccharides present in the polysaccharide extract and the corresponding histogram was plotted accordingly (Fig 7)

### Zebrafish genotoxic experiments

### Zebrafish rearing and exposure

All the methods were approved and performed in accordance with the relevant guidelines and regulations of the Institutional Ethical Committee (IEC) of KIIT University. Zebrafish embryos were obtained from mating adult wild type zebrafish using a 2:1 female: male ratio, and the fertilized embryos were reared in embryo water (0.06% sea salt). Culture density of the fertilized embryos per mating was averaged to 100. The fertilized embryos of 6-hour postfertilization (hpf) were observed under microscope and pipetted in 6-well microplates for the exposure experiments. Zebrafish embryos at 12 hpf (n = 120), in the shield stage, were divided into three groups of 40 embryos each: Control, UVB-exposed, and IOP treated UVB-exposed groups. For UVB exposure and IOP treatment, the embryos (n = 80, 12 hpf) were exposed to UVB dose of 12 J/m^2^/s using a UV cross linker hybridization chamber that emitted a range of 310–312 nm UVB light for 10 secs. Then, the embryos were reared in the embryo water at 28 °C. IOP treatment (2.5 mg/mL) was applied to half of the UVB-exposed embryos (n = 40, 24 hpf) that were labelled IOP-treated-UVB exposed group. The IOP concentration was selected based on our previous study that showed that the adopted dose did not affect the normal development of zebrafish embryos [12]. The embryos were then grown for upto 7 days in the same medium and the fetal embryo toxicity and genotoxicity was assessed. The control group embryos were reared without any treatment in the embryo water for the same duration. The assays were performed in triplicates. Morphological deformities based on phenotypic observations were recorded microscopically during embryonic development. The three zebrafish embryo groups were further assessed for genotoxicity assays using acridine orange staining, alkaline comet assay and qRT-PCR.

### Acridine orange staining

Fluorescent acridine orange dye that binds to DNA and emits green fluorescence was used to stain the all the three groups at 5 dpf for qualitative assessment of the DNA damage and apoptosis. Briefly, control, UVB-exposed and IOP-treated (2.5 mg/mL)-UVB-exposed zebrafish embryos of 5 dpf (n=10/group) were stained with 5 μg/mL acridine orange for 20 min, followed by washing the excess stain with embryo water. The corresponding images were captured in the GFP (green channel) of inverted fluorescent microscope (EVOS, Thermo Fischer Scientific, USA) to assess DNA damage (green dots represented apoptotic cells). The assays were performed in triplicates.

### Alkaline comet assay for DNA damage analysis

DNA fragmentation is considered as one of the endpoints of genotoxicity. Genotoxicity was assessed by evaluating the DNA breaks using the alkaline comet assay and assessing the amount of DNA in the tail region (tail intensity). For this, control zebrafish, UVB-exposed zebrafish, and zebrafish treated with IOP (2.5 mg/mL) after UVB exposure were homogenized thoroughly in a micropestle in two independent experiments at 5 dpf and 7 dpf (n = 5/group). Then, 1 mL Dulbecco’s Modified Eagle Medium (DMEM) with 10% FBS was added, and centrifuged for 5 min at 700 g at 15 °C. The pellet was retrieved and diluted in 1X PBS buffer pH 7.4 to form the cell suspension. Cell homogenate (10 μL) was mixed with 0.8% low melting agarose and were dropped onto a glass slide precoated with normal melting agarose (1.5%), followed by covering with a cover slip. The slides were kept at 4°C for 15 minutes for drying, followed by incubation in a cold lysis solution containing 100 mM EDTA, 2.5M sodium lauryl sulphate, 1% Triton X-100, and 10% DMSO (pH 13.0) in the dark at 4°C for overnight. Following incubation, the slides were dipped in a neutralizing solution containing 400 mM Tris at pH 7.4 for 30 minutes. Then, for assessing the unwinding of the DNA, electrophoresis was performed in a cold alkaline buffer (12 g/L NaOH and 0.37 g/L EDTA, pH 11) at 25 V for 30 minutes. Post electrophoresis, the slides were washed in distilled water, and fixed with 70% ethanol for 5 minutes. Finally, the slides were stained with ethidium bromide (5 mg/mL) for 5 minutes and analyzed by fluorescence microscopy at 4X and 40 X magnifications (fluorescence at emission (500 nm) and excitation (530 nm) with an inbuilt image system (EVOS M5000, Thermo Fisher, USA). The images were analyzed using Image J analysis software for assessing the tail intensity in the three groups. All the assays were conducted in triplicates.

### Gene expression analysis using qRT-PCR

Relative expression of DNA repair genes, *XRCC5, XRCC6, RAD51, GADD45, BAX,* and *P53* were assessed using Syber Green qRT-PCR. Beta actin was used as the reference gene. Briefly, total RNA was extracted from 5 dpf zebrafish larvae from the 3 groups [control, UVB-exposed and IOP-treated (2.5 mg/mL) UVB exposed] by using Trizol and converted to cDNA following standard protocol. cDNA was amplified using a Sybr-green based qPCR master mix and specific primers (Supplementary table 1). The PCR thermal profile consisted of an initial denaturation of at 95 °C for 2 mins, followed by 33 cycles of 30 seconds at 95 °C, 30 seconds at 55-62°C (varied as per primers), and 30 seconds at 72°C, and final extension at 72°C for 3 mins. Qauntification and data analysis of the respective genes were assessed using the CFX manager of real-time PCR (Bio-Rad). The Ct values were calculated for each gene and were compared with the reference gene beta-actin, followed by estimation of ΔΔCt and fold change (2-△△ct) to assess the relative gene expression among the different groups.

### Statistical analysis

The data obtained on the developmental characteristics and alkaline comet assay attribute, tail intensity (%) were reported as the mean ± SD for all experiments independently. The relative gene expression for all the DNA repair genes was estimated with reference to the beta-actin gene in all the three groups. The relative gene expression and the tail intensity of UV exposed zebrafish and IOP-treated UVB exposed group were compared with respect to control zebrafish larvae using one-way analysis of variance (ANOVA), followed by multiple comparison tests in GraphPad Prism software. A p < 0.05 was considered statistically significant for assessing the association of the variables in context to genotoxicity.

## Ethical statement

All the methods were approved and carried out in accordance with relevant guidelines and regulations of the Institutional Ethical Committee (IEC) of KIIT University.

## Author Contributions

JIE conceived the experiment design, performed experiments and data analysis and was involved in the manuscript drafting. BD conceived performed and designed the experiments and also involved in writing and editing the manuscript. Both authors reviewed and discussed their views on the manuscript.

